# *Lysinibacillus fusiformis* M5 induces increased complexity in *Bacillus subtilis* 168 colony biofilms via hypoxanthine

**DOI:** 10.1101/118125

**Authors:** Ramses Gallegos-Monterrosa, Stefanie Kankel, Sebastian Götze, Robert Barnett, Pierre Stallforth, Ákos T. Kovács

## Abstract

In recent years, biofilms have become a central subject of research in the fields of microbiology, medicine, agriculture, or systems biology amongst others. The sociomicrobiology of multispecies biofilms, however, is still poorly understood. Here, we report a screening system that allowed us to identify soil bacteria, which induce architectural changes in biofilm colonies when cocultured with *B. subtilis*. We identified the soil bacterium *Lysinibacillus fusiformis* M5 as inducer of wrinkle-formation in *B. subtilis* colonies mediated by a diffusible signaling molecule. This compound was isolated by bioassay-guided chromatographic fractionation. The elicitor was identified to be the purine hypoxanthine using mass spectrometry and nuclear magnetic resonance (NMR) spectroscopy. We show that the induction of wrinkle formation by hypoxanthine is not dependent on signal recognition by the histidine kinases KinA, KinB, KinC, and KinD, which are generally involved in phosphorylation of the master regulator Spo0A. Likewise, we show that hypoxanthine signaling does not induce the expression of biofilm-matrix related operons *epsA-O* and *tasA-sipW-tapA*. Finally, we demonstrate that the purine permease PbuO, but not PbuG, is necessary for hypoxanthine to induce an increase in wrinkle formation of *B. subtilis* biofilm colonies. Our results suggest that hypoxanthine-stimulated wrinkle development is not due to a direct induction of biofilm-related gene expression, but rather caused by the excess of hypoxanthine within *B. subtilis* cells, which may lead to cell stress and death.

**IMPORTANCE:** Biofilms are a bacterial lifestyle with high relevance regarding diverse human activities. Biofilms can be favorable, for instance in crop protection. In nature, biofilms are commonly found as multispecies communities displaying complex social behaviors and characteristics. The study of interspecies interactions will thus lead to a better understanding and use of biofilms as they occur outside laboratory conditions. Here, we present a screening method suitable for the identification of multispecies interactions, and showcase *L. fusiformis* as a soil bacterium that is able to live alongside *B. subtilis* and modify the architecture of its biofilms.

## INTRODUCTION

Biofilms are microbial populations formed by cells living in high density communities attached to biotic or abiotic surfaces. These cells are often encased in a matrix of polymeric substances that provide the whole population with an increased resistance against environmental stress (1, 2). Furthermore, these communities exhibit highly complex structural organization and social behavior. Thus, biofilms have become an increasingly studied research subject by microbiologists, especially when it became apparent that this lifestyle is widely spread among bacteria and involved in a multitude of biological processes (3, 4). Although much attention has been given to medically relevant biofilms (5, 6), scientists have also studied biofilms in the context of industrial applications (7), bioremediation (8), and crop protection (9).

In nature, biofilms rarely occur as single-species populations, but rather as mixed communities of diverse bacteria and other microorganisms. This leads to complex interactions between the different members of the community, usually involving communication networks based on chemical signals (10). Additionally, microorganisms need to sense and efficiently adapt to a wide array of environmental cues in order to efficiently regulate biofilm formation (11).

*Bacillus subtilis* is a soil-dwelling Gram-positive bacterium that has become a model for biofilm research. On agar plates, *B. subtilis* can form large colonies with remarkably complex architecture, while in liquid medium it forms robust floating biofilms known as pellicles. Both forms of biofilms are characterized by a wrinkled surface, which has been associated to the production of exopolysaccharides, biofilm maturation, and mechanical forces concomitant with an increased population complexity (12–14). Moreover, these biofilms display intricate cell heterogeneity, i.e. some cells become matrix producers, while others either produce exoenzymes to harvest nutrients, or form resistant structures known as spores (15, 16). The development of this population heterogeneity is regulated by a complex gene regulatory network involving various sensing kinases i.e. Kin kinases, DegS, and ComP, and their concomitant response regulators: Spo0A, DegU, and ComA respectively; and other downstream regulators, e.g. SinI and SinR (15, 17, 18).

Biofilms produced by *B. subtilis* are not only a good research model, they are also currently applied in crop protection (19, 20), and spores of this organism are readily commercialized as a biocontrol agent for agriculture. *B. subtilis* is a prolific producer of secondary metabolites and many potent antimicrobial compounds inhibiting both bacteria and fungi have been identified (21–23). In addition, it has also been shown that *B. subtilis* activates biofilm-related gene expression in response to chemicals produced by other bacteria closely related to it, for instance by other members of the *Bacillus* genus (24). Interestingly, the signaling role of the molecules can be independent from other effects of the compounds, as in the case of antimicrobial thiazolyl peptides, which can induce biofilm-matrix production in *B. subtilis* even when separated from their antibiotic activity (25). Moreover, other organisms such as *Pseudomonas protegens*, are able to inhibit cell differentiation and biofilm gene expression in *B. subtilis*, possibly as a competition strategy during root colonization (26).

*B. subtilis* successfully inhabits a congested and competitive ecological niche (27–29), and it is to be expected that this organism has a finely tuned regulatory network governing community behavior. Therefore, further study of the signaling mechanisms that influence *B. subtilis* biofilm formation may enhance the use of this organism, both in biotechnological applications as well as a research model. However, the identification of signals that induce biofilm formation is a poorly investigated field of study, possibly due to the greater general interest in the removal of biofilms in various medical and industrial settings (5, 30–32). Thus, we have established a co-cultivation-based screening method to identify signaling molecules that promote the development of wrinkles in colony biofilms of *B. subtilis*. Using this system, we identified ecologically relevant soil bacteria that are able to induce the formation of large wrinkles in colony biofilms of *B. subtilis*. The majority of these bacteria are members of the family *Bacillaceae*, to which *B. subtilis* belongs. The strain with the clearest wrinkle-inducing effect was identified as *Lysinibacillus fusiformis* M5. The observed effect on *B. subtilis* is dependent on a diffusible signaling molecule, which was identified as hypoxanthine using bioassay-guided fractionation and subsequent structure elucidation using various spectroscopic and spectrometric methods. The induction of wrinkles by hypoxanthine was not dependent on Kin kinases signal transduction, and the expression levels of operons responsible for the production of biofilm matrix components, *epsA-O* and *tapA-sipW-tasA*, remained unaffected. We show that uptake of hypoxanthine by permease PbuO is necessary for the increased induction of wrinkle formation in *B. subtilis* biofilm colonies. We therefore suggest that hypoxanthine induces the formation of wrinkles by introducing a metabolic change in *B. subtilis* cells, rather than by direct stimulation of biofilm-related gene expression.

## RESULTS

### Screening of soil bacteria that induce structural changes in colonies of *B. subtilis*

We screened a collection of 242 strains isolated from two distinct soil sampling sites in Mexico, in order to identify bacteria that are able to induce biofilm formation or increased complex colony architecture of *B. subtilis*. Importantly, our assay aimed to discover alterations in biofilm colony architecture that was different from the previously described method that identified soil-derived microbes that induce gene expression related to biofilm formation of *B. subtilis* (24). While some *B. subtilis* strains, such as *B. subtilis* NCIB 3610, easily and spontaneously form biofilms, we used a strain that would form architecturally complex colonies only in the presence of specific inducers or nutrient rich conditions. Thus, even weak biofilm-inducing effects would not be overseen. We used *B. subtilis* 168 (Jena stock), a strain that can only form architecturally complex colonies when grown on glucose-rich medium or exposed to signaling molecules as those present in plant root exudates (33). The strain of *B. subtilis* used for the assay (TB48) carried a P_hyperspank_-mKATE reporter fusion in order to facilitate the identification of *B. subtilis* from the soil strains in mixed colonies. The reporter strain was mixed in different ratios with the bacterial isolates and allowed to form colonies on 2×SG medium for 72 hours. Single strain colonies of *B. subtilis* and the soil-derived isolates were also grown as neighboring colonies under the same conditions, inoculated with a spatial distance of 5 mm between each other to examine their interactions. The majority of these interactions resulted in the apparent killing of one strain by the other, producing a colony identical to the pure culture colony of the surviving partner (data not shown). However, 36 soil strains were able to grow alongside *B. subtilis*, mainly by creating a colony where the strains segregate in sectors. Interestingly, the *B. subtilis* sectors of these mixed colonies showed an increased architectural complexity by forming large wrinkles and a rugose colony surface, compared to its pure colonies, which remained flat (Fig. 1). Furthermore, when single strain colonies of these soil-derived isolates were grown close to *B. subtilis* colonies, the induction of wrinkle formation could be observed in the areas of the *B. subtilis* colony that are closest to the soil strain, but not in the other regions of the colony (Fig. 1). These results suggest that the aforementioned bacteria produce signals that can induce an increased architectural complexity in *B. subtilis* colony biofilms.

**Figure 1.**
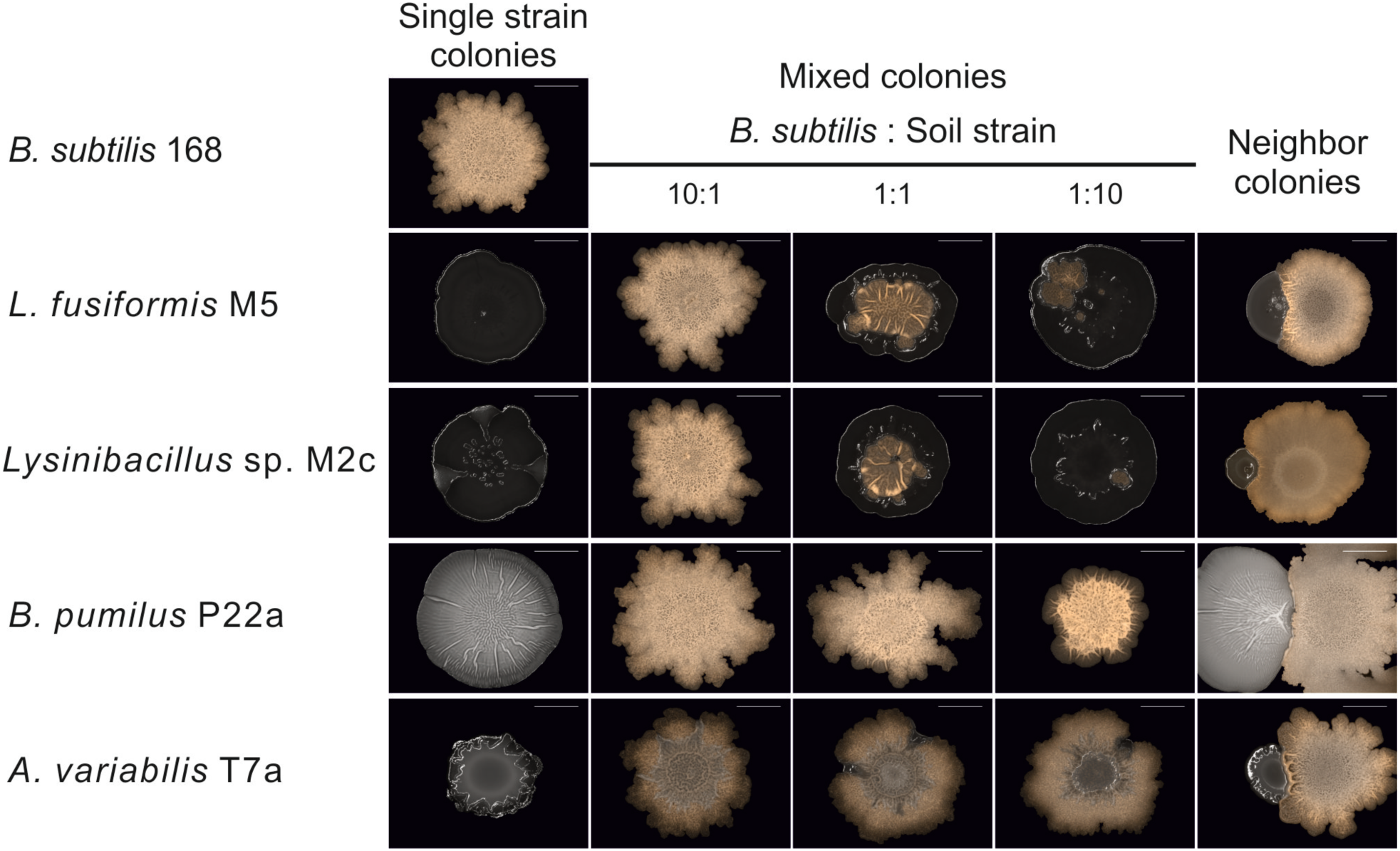
Single strain and mixed colonies of *B. subtilis* and selected soil strains. *B. subtilis* was differentiated from the soil strains using fluorescence emission (false-colored orange) from a reporter expressed by the P_*hyperspank*_-mKATE construct. Colonies are shown after 72 h of incubation. Neighbor colonies were inoculated at 5 mm from each other. Scale bars represent 5 mm.

### Structural changes in *B. subtilis* colonies are induced by diffusible signals produced by soil bacteria

In order to elucidate if the observed induction of wrinkle formation is caused by a diffusible signal molecule or due to direct cell-cell interactions, we designed an assay to test the cell-free supernatants of the selected soil strains. In this assay, we used cotton discs infused with cell-free supernatant to simulate colonies of the tested soil strains. *B. subtilis* was inoculated at a distance of 5 mm next to the cotton discs and allowed to grow for 72 hours. Over this period, the growing colony surrounded the cotton discs, coming into contact with the freely diffusing compounds in the supernatants.

Using this assay, we observed that the cell-free supernatants of four bacterial isolates were able to induce efficiently the formation of wrinkles in the adjacent *B. subtilis* colony. Importantly, this phenomenon was observed in the periphery of the cotton discs, but not in the areas of the colony farther away from it (Fig. 2). We note that neither the cell-free supernatant of *B. subtilis* itself, nor the medium used to obtain the supernatants, showed the capacity to induce increased wrinkle formation in *B. subtilis* colonies under our tested conditions (Fig. 2). We concluded that the induction of wrinkle formation was due to a diffusible signaling molecule produced by these soil organisms.

**Figure 2.**
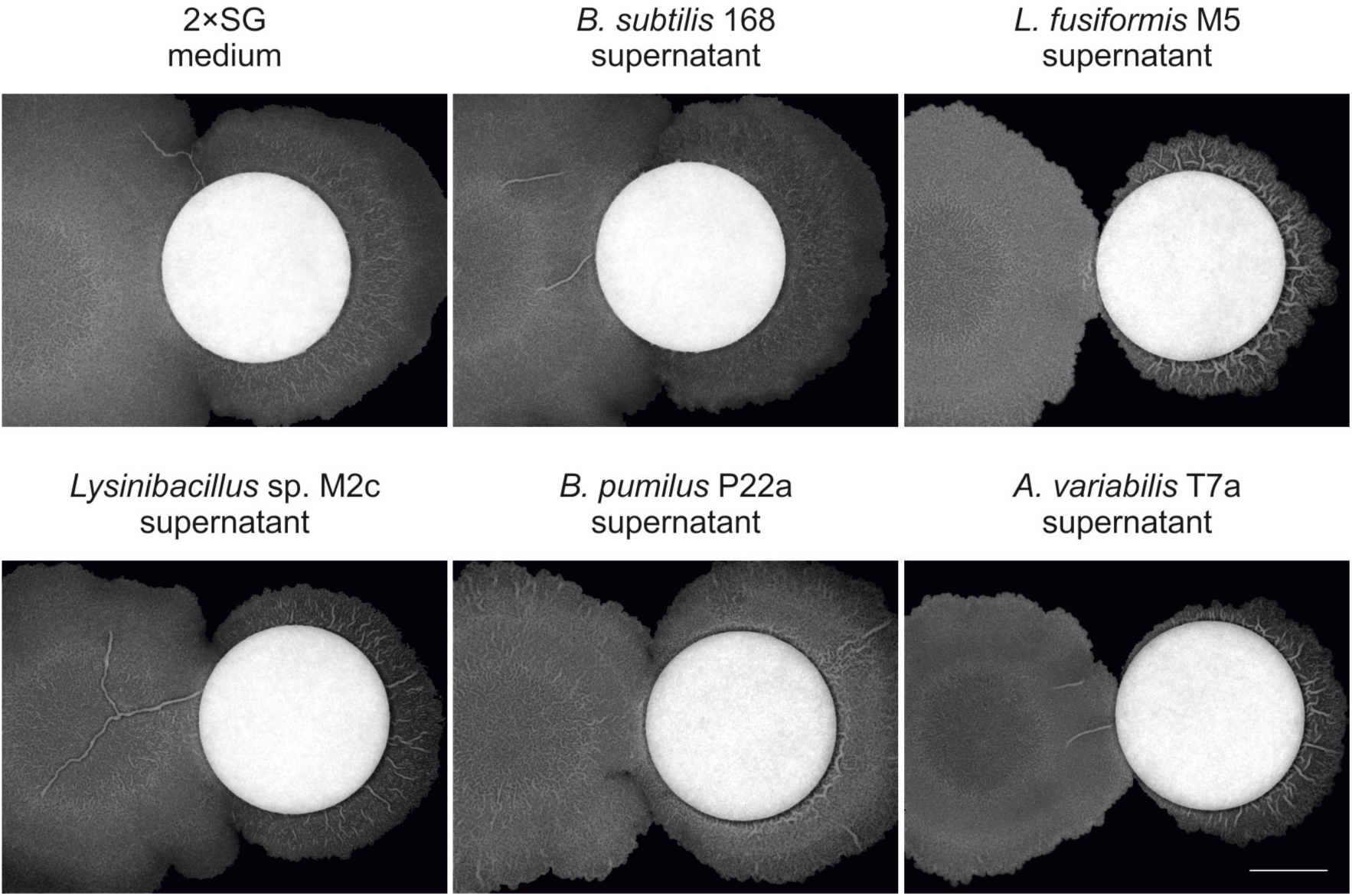
Effect of cell-free supernatants of soil strains on the development of *B. subtilis* 168 biofilm colonies. Colonies were inoculated with 2 µL of culture at 5 mm from white cotton discs impregnated with 50 µL of cell-free supernatant of soil bacteria or 2×SG medium. The discs were reimpregnated with 25 µl of the corresponding medium or supernatant every 24 h. Bright-field images of colonies are shown after 72 h of incubation. The scale bar represents 5 mm.

Using 16S rRNA locus sequencing, we characterized those soil strains whose supernatant could best stimulate wrinkle formation in *B. subtilis*. The majority of the sequenced strains were found to be members of the same phylogenetic family as *B. subtilis*, such as *Bacillus pumilus* or *L. fusiformis*. The only exception was a strain identified as the γ-proteobacterium *Acinetobacter variabilis*.

### Hypoxanthine identified in the supernatant is responsible for wrinkle induction

The strongest induction of wrinkle formation (defined as the appearance of tall wrinkles and a rugose colony surface) was observed with the supernatant of the soil derived strain identified as *L. fusiformis* M5. For this reason, we decided to further investigate the respective signaling molecule produced by this bacterium using a wrinkle formation assay. Bio-assay-guided fractionation allowed us to identify a compound from *L. fusiformis* M5 that induced a similar phenotype as observed when *B. subtilis* and *L. fusiformis* M5 were co-cultured. To this end, supernatant of *L. fusiformis* M5 was lyophilized and fractionated using Sephadex G20 as stationary phase. Each fraction was applied to cotton discs and placed on an agar plate in the vicinity of *B. subtilis.* The fraction that induced wrinkle formation was sub-fractionated by high-performance liquid chromatography (HPLC) using a hypercarb column as stationary phase. Repeating this procedure led to the isolation of a homogeneous compound whose structure was subsequently elucidated via a combination of nuclear magnetic resonance (NMR) spectroscopy and high-resolution mass spectrometry (HR-MS). Finally, hypoxanthine was identified as the inducer of wrinkle formation (Fig. 3). We further validated our findings using commercial hypoxanthine, which showed the same retention time as the isolated hypoxanthine (Fig. 3).

**Figure 3.**
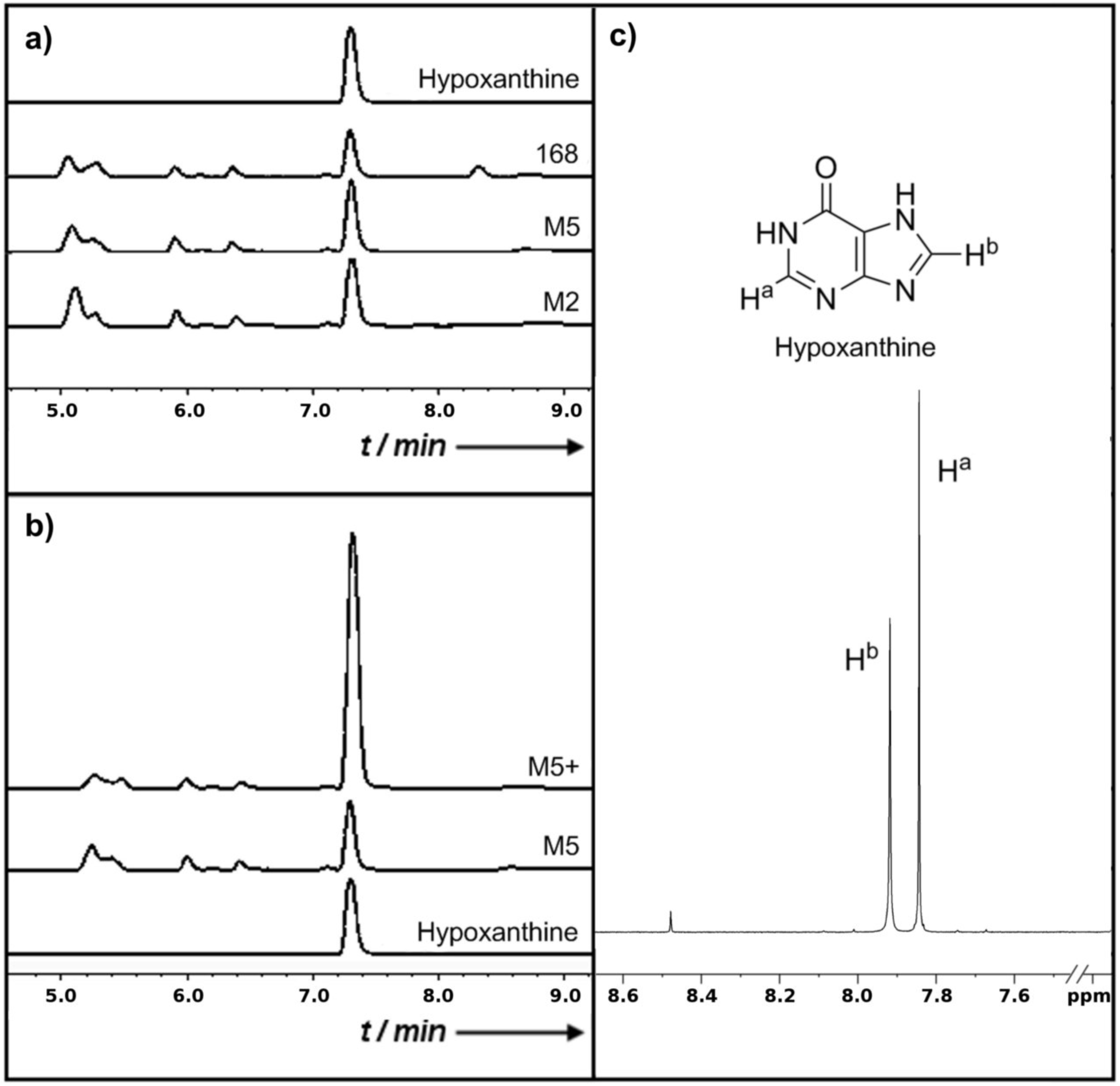
Characterization of hypoxanthine standard and cell-free supernatants of *B. subtilis* 168 (168), and soil isolates *L. fusiformis* M5 (M5) and *Lysinibacillus* sp. M2c (M2). a) HPLC chromatograms of cell-free supernatants compared to a standard solution of hypoxanthine. b) HPLC chromatogram of *L. fusiformis* M5 cell-free supernatant compared with a standard solution of hypoxanthine, and the *L. fusiformis* M5 cell-free supernatant spiked with a standard of hypoxanthine (M5+). c) ^1^H NMR spectrum of isolated hypoxanthine (DMSO-d6, 600 MHz).

We used the colony wrinkle formation assay to test if hypoxanthine (as a 25 mM solution in 0.05 N NaOH) alone can induce the formation of wrinkles in *B. subtilis* colony biofilms. In addition, guanine and xanthine were also tested using this methodology, since these purines can be found in the same metabolic pathways as hypoxanthine in *B. subtilis*. We found that hypoxanthine and guanine were able to induce the formation of tall wrinkles in *B. subtilis* colonies, while xanthine could not (Fig. 4a). Interestingly, guanine can be deaminated during purine catabolism to produce hypoxanthine, which can then be oxidized to produce xanthine (Fig. 4b). These results suggest that a metabolite derived from guanine or hypoxanthine, but not xanthine, may be responsible for the observed formation of wrinkles.

**Figure 4.**
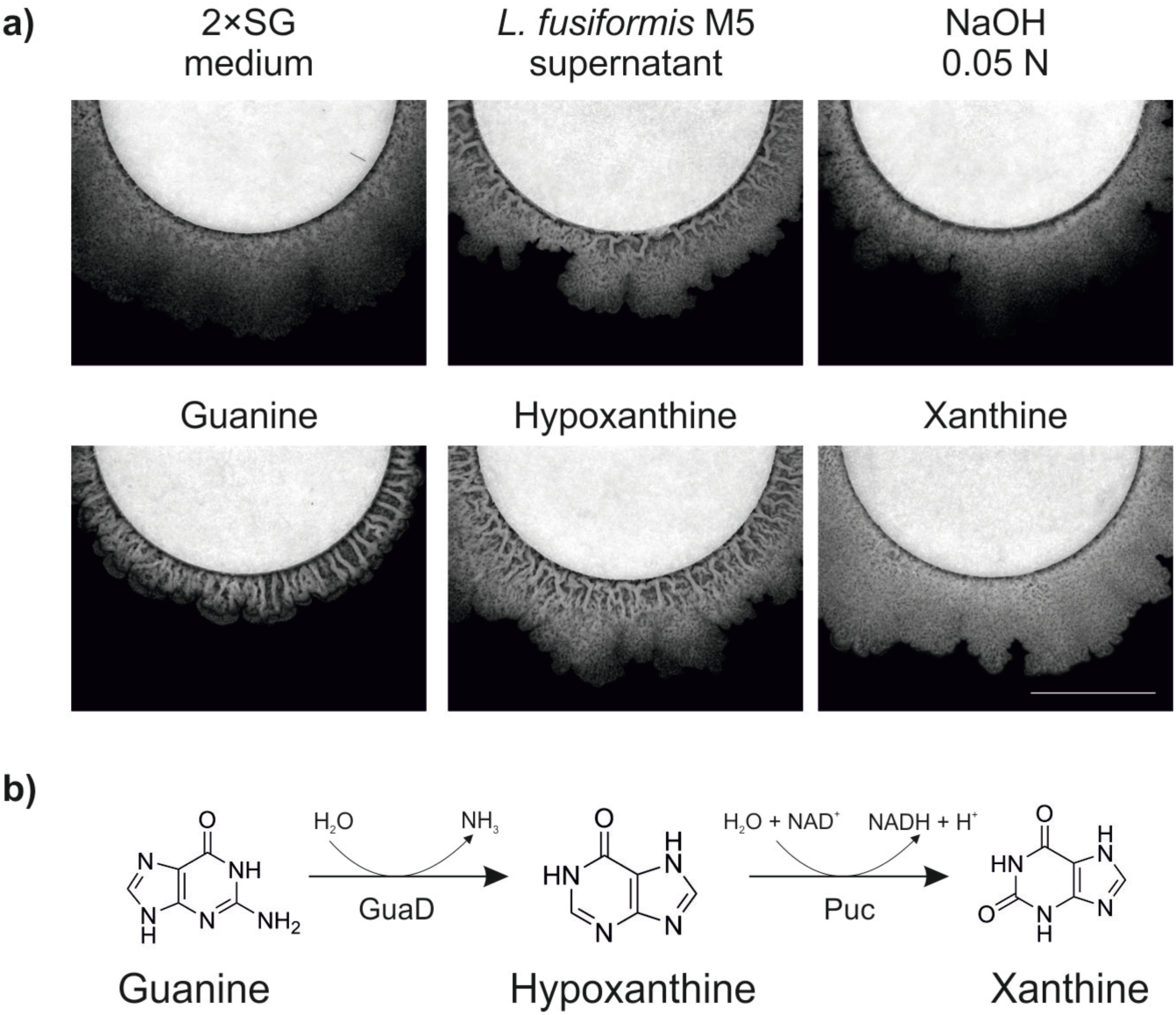
Effect of cell-free supernatant of *L. fusiformis* M5, guanine, hypoxanthine, and xanthine on the development of biofilm colonies of *B. subtilis* 168 (a) and catabolic pathway of guanine, hypoxanthine and xanthine (b). GuaD: guanine deaminase. Puc: hypoxanthine/xanthine dehydrogenases (PucA-E). Bright-field images of colony areas adjacent to the cotton discs are shown after 72 h of incubation. The scale bar represents 5 mm.

### Hypoxanthine signaling is not mediated by the activity of individual Kin kinases

In *B. subtilis*, the transcriptional regulator Spo0A controls the expression of several biofilm-related operons, including those responsible for the production of the exopolysaccharide and protein components of the biofilm matrix (*epsA-O* and *tapA-sipW-tasA*, respectively) (17). Five sensor kinases (KinA, KinB, KinC, KinD, and KinE) have been identified in *B. subtilis*, and four of them (KinA-D) are known to activate Spo0A via a phosphorelay depending on environmental signals (11, 15, 34). We wanted to determine if any of these kinases is involved in hypoxanthine-mediated induction of wrinkles. Therefore, we compared the effect that the supernatant of *L. fusiformis* M5 has on kinase-mutant strains of *B. subtilis* using the colony wrinkle formation assay. We expected that, should one of these kinases be responsible for sensing hypoxanthine, the corresponding mutant strain would no longer show increased induction of wrinkle formation. Interestingly, all the mutant strains were still able to develop highly wrinkled colonies when exposed to the supernatant, when compared to the corresponding colonies exposed to 2×SG medium (Fig. 5). This result suggests that hypoxanthine-mediated induction of increased architectural complexity is not mediated by the activation of Spo0A via a single Kin kinase. We note that more than one Kin kinase could be responsible for detecting hypoxanthine, in which case the deletion of a single *kin* kinase gene would not prevent *B. subtilis* to form wrinkled colonies when exposed to hypoxanthine.

**Figure 5.**
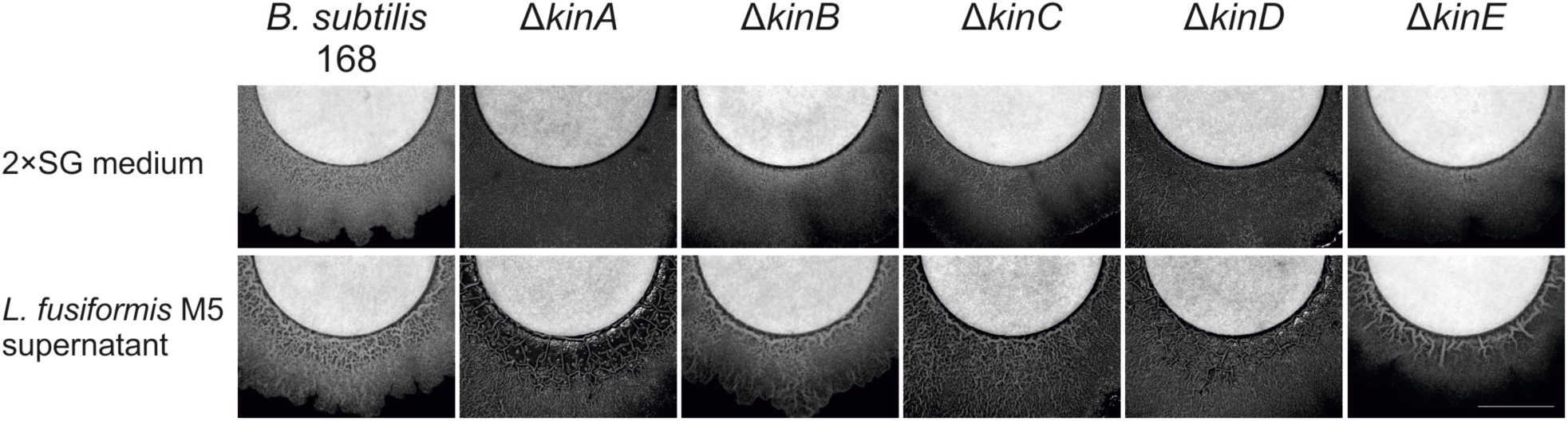
Effect of cell-free supernatant of *L. fusiformis* M5 on the development of biofilm colonies of *B. subtilis* 168 and knock-out mutants of kin-kinase genes. Colonies were inoculated with 2 µl of culture at 5 mm from white cotton discs impregnated with 50 µL of cell-free supernatant of *L. fusiformis* M5 or 2×SG medium. The discs were reimpregnated with 25 µL of the corresponding medium or supernatant every 24 h. Bright-field images of colony areas adjacent to the cotton discs are shown after 72 h of incubation. The scale bar represents 5 mm.

### Expression levels of genes responsible for biofilm matrix production are not affected by hypoxanthine signaling

The *epsA-O* and *tapA-sipW-tasA* operons are related to the production of the exopolysaccharide and protein components of the *B. subtilis* biofilm matrix, respectively. Changes in the expression levels of these operons are associated to a maturating biofilm, and show spatiotemporal variation during its development (16, 35, 36). To further examine if biofilm matrix-related genes are involved in the induction of wrinkles by hypoxanthine, we monitored the expression of P_*epsA*_-*gfp* and P_*tapA*_-*gfp* fluorescent reporter fusions in colonies of *B. subtilis* using the colony wrinkle formation assay. Fluorescence emission was examined only in the sections of the colonies directly adjacent to the cotton discs at 3 time points: (i) when *B. subtilis* has encircled the discs (40 hours), (ii) when the colony started to expand from the disc and showed the onset of wrinkle formation (50 hours), and (iii) when the colony has developed wrinkles and expanded (65 hours). The examined strains also carried a P_*hyperspank*_-*mKATE* reporter fusion to adjust for colony growth. Under these conditions, the expression from P_*epsA*_ and P_*tapA*_ in these colonies showed no statistical differences when exposed to cotton discs infused with 2×SG medium or supernatant of *B. subtilis,* as compared to those infused with supernatant of *L. fusiformis* M5 (one-way ANOVA: P < 0.05, n = 4-8 independent colonies) (Fig. 6).

**Figure 6.**
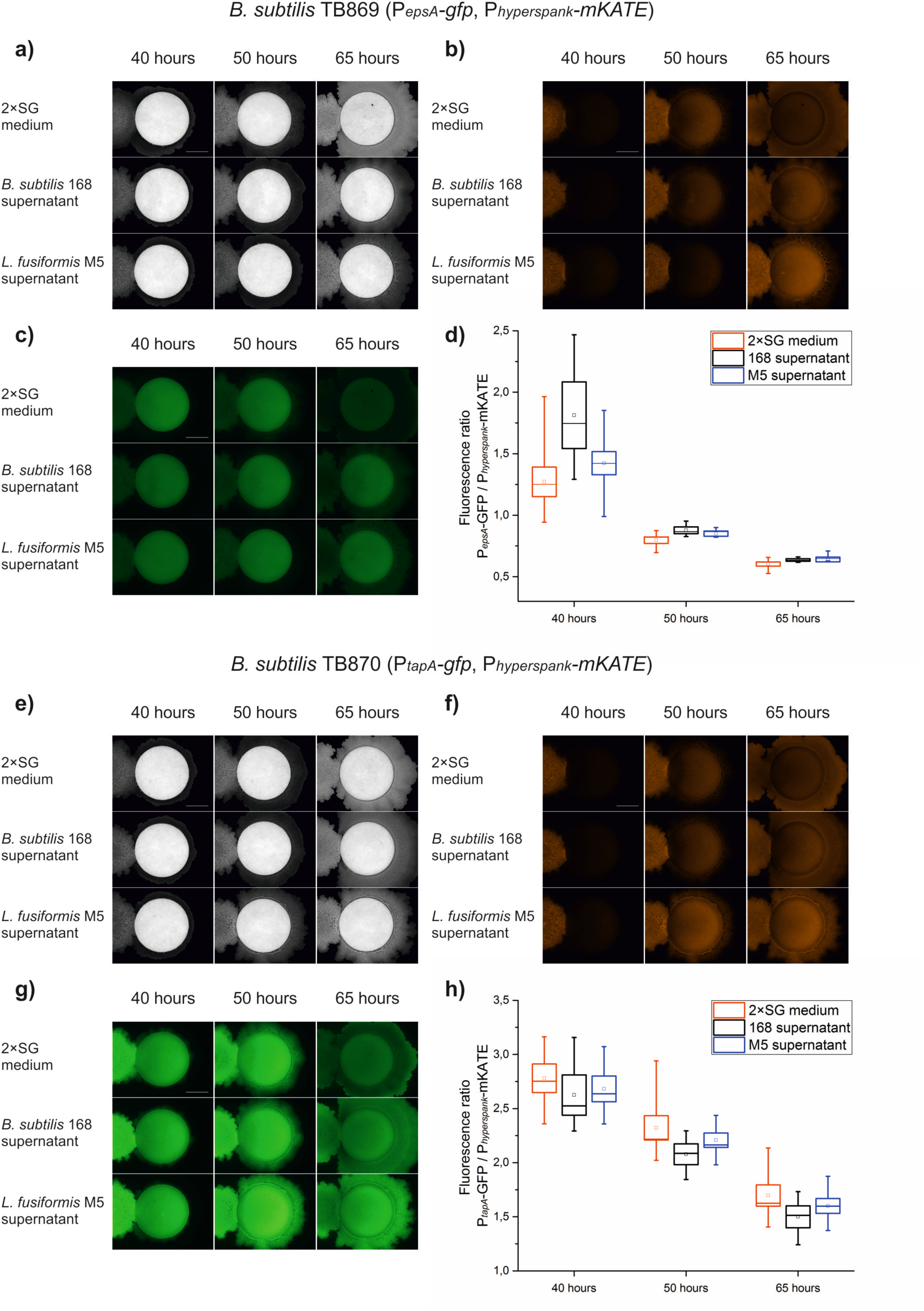
Comparison of fluorescence emission of *B. subtilis* strains carrying the constitutive P_*hyperspank*_-*mKATE*, and the P_*epsA*_-*gfp* (a-d) or P_*tapA*_-*gfp* (e-h), reporter fusions. Bright field (a and e), red fluorescence (b and f), and green fluorescence images (c and g) of representative *B. subtilis* biofilm colonies exposed to cell-free supernatants of *B. subtilis* 168, *L. fusiformis* M5 and 2×SG medium. Box plots of the ratio of green and red fluorescence emission of biofilm colonies of *B. subtilis* TB869 (d) and TB870 (h) exposed to cell-free supernatants of *B. subtilis* 168, *L. fusiformis* M5 and 2×SG medium at different time points. Scale bars represent 5 mm. Box plots (d and h) represent fluorescence ratios of at least 4 independent colonies, processed as described in material and methods. (one-way ANOVA: P < 0.05, n = 4-8 independent colonies).

We decided to further test if the products of the *epsA-O* and *tapA-sipW-tasA* operons are necessary for the observed development of wrinkles. We used the wrinkle formation assay to test the effect of the supernatant of *L. fusiformis* M5 on mutant strains of *B. subtilis* that are unable to produce the exopolysaccharide (Δ*epsA-O*) or protein (Δ*tasA*) matrix component. After 72 hours of incubation, the tested *B. subtilis* strains expanded and surrounded the infused cotton discs, but were unable to develop wrinkles and showed a flat and mucoid colony surface (Fig. 7).

**Figure 7.**
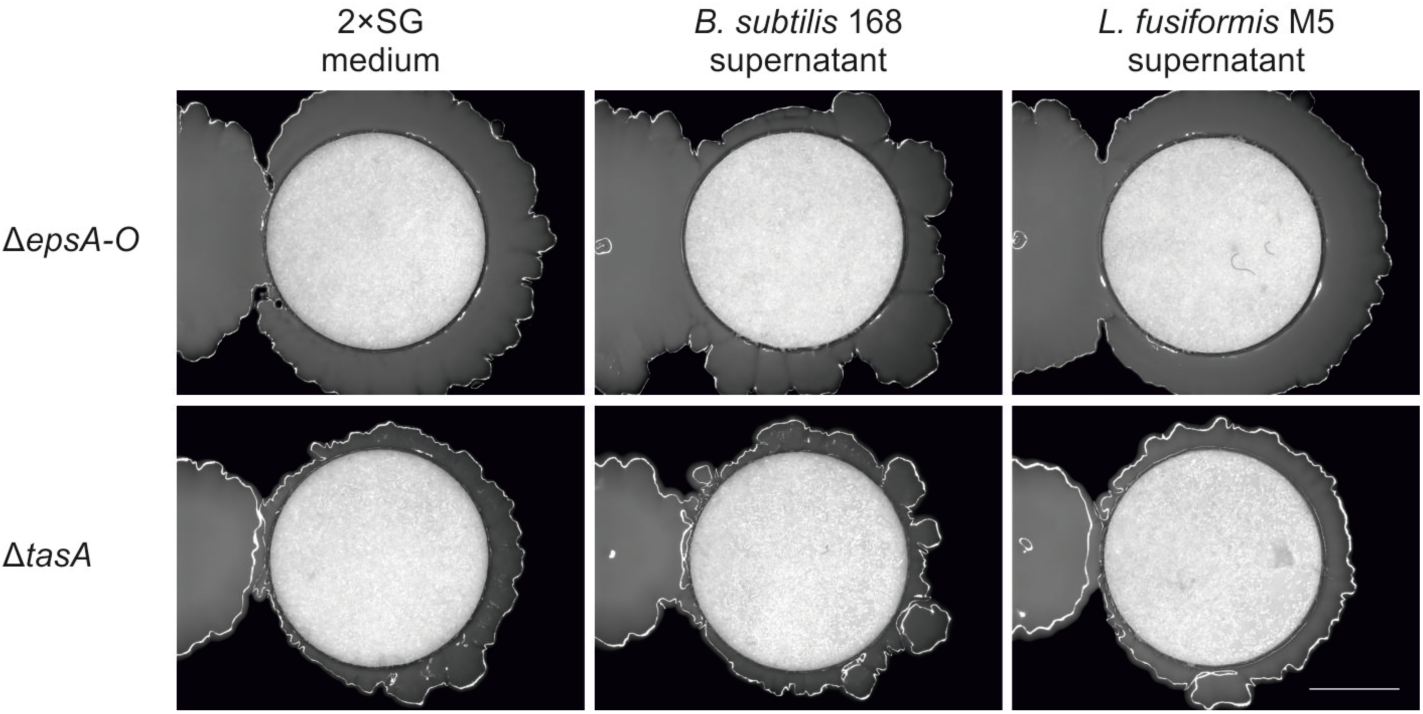
Effect of cell-free supernatant of *L. fusiformis* M5 on the development of colonies of *B. subtilis* 168 knock-out mutants of biofilm matrix biosynthetic operon *epsA-O* and *tasA* gene. Colonies were inoculated with 2 µL of culture at 5 mm from white cotton discs impregnated with 50 µL of cell-free supernatants of bacterial culture or 2×SG medium. The discs were reimpregnated with 25 µL of the corresponding medium or supernatant every 24 h. Bright-field images of colony areas adjacent to the cotton discs are shown after 72 h of incubation. The scale bar represents 5 mm.

Taken together, these results suggest that the increased colony wrinkle formation induced by hypoxanthine is not directly associated with a large increased in expression from the biofilm matrix operons; however, the production of a biofilm matrix is necessary for the development of wrinkles.

### Cell death correlates with wrinkle formation

It has been shown previously that localized cell death can be a trigger for wrinkle formation in biofilm colonies. This happens as a consequence of mechanical forces converging on zones of cell death, which lead to a buckling of the biofilm and the rise of tall wrinkles (13). Based on our previous results, we hypothesized that hypoxanthine, or a metabolite formed during its catabolism, may cause cell death in *B. subtilis*. This would produce mechanical stress in the developing biofilm and lead to buckling and wrinkle formation. Thus, we used Sytox Green to assess the distribution of dead cells in the colony wrinkle formation assay. Sytox Green is a commercially available fluorescent nucleic acid stain that has been established as a reporter of cell death for bacteria (37). For this assay, we used a *B. subtilis* strain that carries a P_*hyperspank*_-mKATE reporter fusion (TB48) in order to facilitate the identification of *B. subtilis* cells that are metabolically active from those that are readily stained by Sytox Green. After 72 hours of incubation, we examined thin cross-sections of colonies that were exposed to 2×SG medium or supernatant from *L. fusiformis* M5. The examined cross-sections corresponded to areas of the colonies adjacent to the cotton discs (Fig. 8a and f). We observed that cell death is localized at the bottom of the biofilm, both on those exposed to 2×SG medium or *L. fusiformis* M5 supernatant (Fig. 8). However, in the cross-sections obtained from the flat colonies of *B. subtilis* exposed to 2×SG medium, the dead cells appear as thin layer of similar width along the length of the cross-section (Fig. 8b-e). In contrast, the cross sections from wrinkled colonies exposed to *L. fusiformis* M5 supernatant revealed aggregates of dead cells that correlate with the wrinkles seen through the colony (Fig. 8g-j). Furthermore, we compared the average green fluorescence of the colony cross-sections (produced by cells stained with Sytox Green) with their own red fluorescence (produced by cells expressing the P_*hyperspank*_-mKATE reporter fusion). We found that the average ratios of green/red fluorescence were significantly higher in the cross-sections of colonies exposed to *L. fusiformis* M5 supernatant (0.69 AU ± 0.10 (standard deviation)), than those of cross-sections from colonies exposed to 2×SG medium (0.33 AU ± 0.02 (standard deviation) (one-way ANOVA: P > 0.05, n = 3 cross-sections from independent colonies). These results confirm that the formation of wrinkles is facilitated by cell death, a phenomenon observed in the presence of the *L. fusiformis* M5 supernatant.

**Figure 8.**
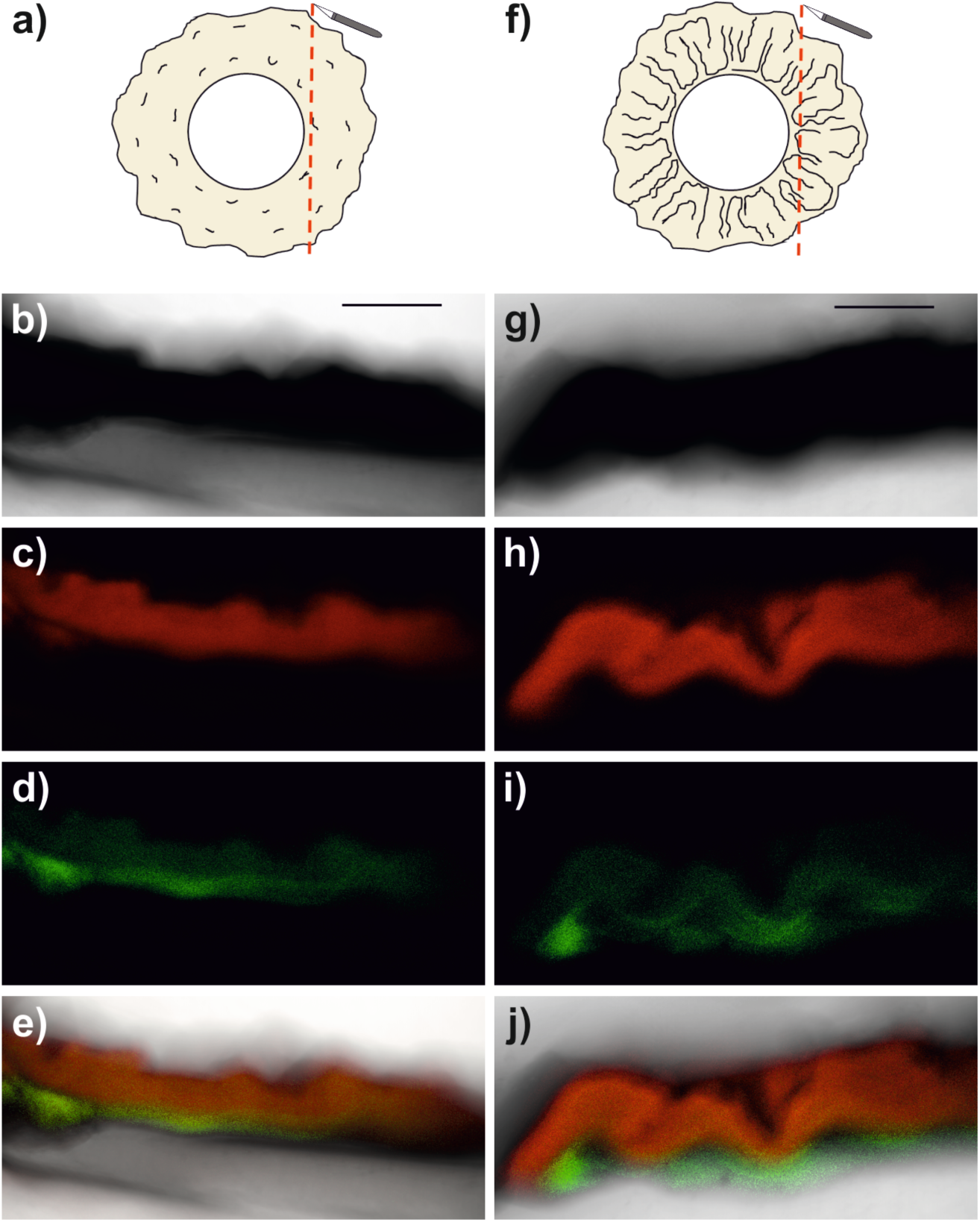
Detection of localized cell death in biofilm colonies of *B. subtilis* exposed to 2×SG medium (b-e) or cell-free supernatant of *L. fusiformis* M5 (g-j). Schematic representations of the cross-sections are shown from areas adjacent to cotton discs (a and b). Transmitted light (b and g), red fluorescence (c and h), green fluorescence (d and i), and composite images (e and j) of colonies of a *B. subtilis* strain carrying the constitutive P_*hyperspank*_-*mKATE* reporter fusion are shown after 72 h of growth on plates with 0.25 µM of Sytox Green. The brightness and contrasts of the images have been enhanced to facilitate the appreciation of fluorescence signals and colony wrinkles. The scale bars represent 250 µm.

### The permease PbuO is necessary for hypoxanthine-induced development of wrinkles

We hypothesized that the observed induction of wrinkle formation may be due to metabolic effects on *B. subtilis* cells derived from an excess of available hypoxanthine provided by the culture supernatant of *L. fusiformis* M5, rather than to direct signal-dependent expression of biofilm-related genes. In this case, hypoxanthine alone would be sufficient to induce increased wrinkle formation, and its uptake by *B. subtilis* would be necessary for this phenomenon.

Using the colony wrinkle formation assay, we observed that different concentrations of hypoxanthine (as solution in 0.05 N NaOH) were able to induce the formation of wrinkles in *B. subtilis* colonies as efficiently as the supernatant of *L. fusiformis* M5. To test whether hypoxanthine internalization is required for the observed wrinkle induction in *B. subtilis*, we analyzed mutant strains of *B. subtilis* that lack *pbuG* or *pbuO*. PbuG is a previously described hypoxanthine/guanine permease (38), while PbuO is a protein paralogous to PbuG annotated as a putative purine permease (see SubtiWiki: http://subtiwiki.uni-goettingen.de/index.php) (39). The hypoxanthine inducing effect disappeared when *pbuO* alone, or in combination with *pbuG,* was deleted (Fig. 9). These final results demonstrated that hypoxanthine uptake is important for induction of wrinkle formation and PbuO is mainly responsible for the observed effect under our experimental conditions.

**Figure 9.**
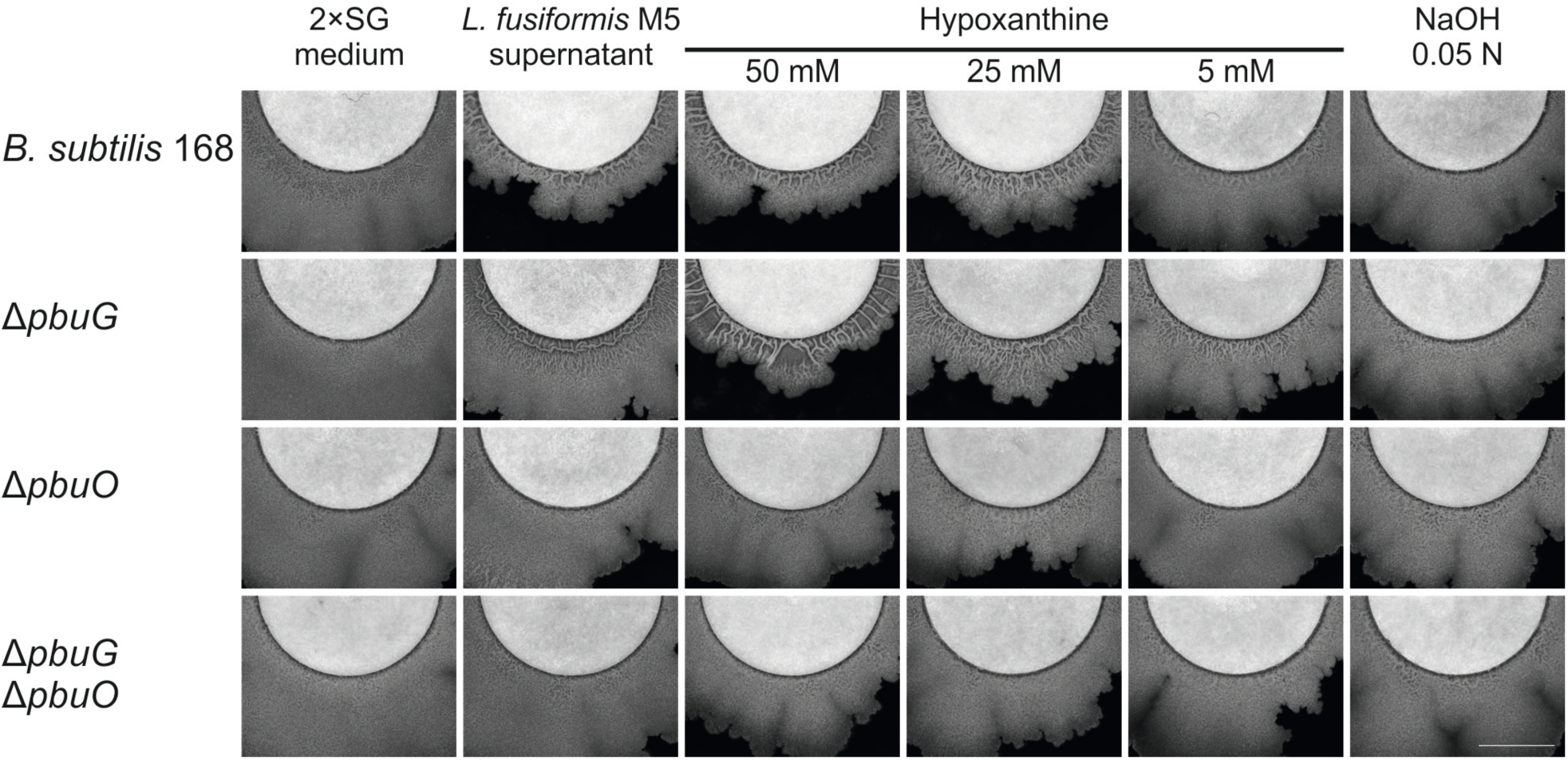
Effect of hypoxanthine and cell-free supernatant of *L. fusiformis* M5 on the development of biofilm colonies of *B. subtilis* 168 and knock-out mutants of *pbuO* and *pbuG* permease genes. Colonies were inoculated with 2 µL of culture at 5 mm from white cotton discs impregnated with 50 µL cell-free supernatant of *L. fusiformis* M5 or 2×SG medium. The discs were reimpregnated with 25 µL of the corresponding medium or supernatant every 24 h. Bright-field images of colony areas close to the cotton discs are shown after 72 h of incubation. The scale bar represents 5 mm.

## DISCUSSION

In this work, we identified a chemical sensing mechanism between *B. subtilis* and other soil bacteria that promotes architectural complexity in colony biofilms. We devised a screening system that allowed us to analyze a collection of soil bacteria, selecting those that could form stable multispecies communities with *B. subtilis*. Using this screening system, we identified *L. fusiformis* M5 as a bacterium capable of inducing an increase of colony wrinkle formation in *B. subtilis* via hypoxanthine as chemical cue.

Hypoxanthine is a purine that plays an important role in the pentose phosphate salvage pathway, which is a mechanism for cells to interconvert nucleosides and nucleobases according to their metabolic needs (40, 41). In *B. subtilis*, hypoxanthine is particularly relevant in this pathway because it can be taken up by cells and used as a substrate by phosphoribosyl-transferases in order to produce inosine monophosphate (IMP), which in turn is converted to adenosine monophosphate or guanine monophosphate (38) (also see SubtiWiki Pathways: http://subtiwiki.uni-goettingen.de/apps/pathway.php?pathway=2) (39). The role of hypoxanthine in eukaryotic cell metabolism has been extensively investigated. In humans, it has been studied in the context of diseases such as gout, Lesch–Nyhan disease, and endothelial cell injury of cardiovascular diseases. Although these conditions have different etiologies and clinical evolution, they have in common an excessive accumulation of hypoxanthine and uric acid, whose catabolism leads to oxidative-stress-induced apoptosis (42–44). In bacteria, hypoxanthine has been mainly studied in relation to DNA damage and mutagenesis due to spontaneous deamination of adenine, which yields hypoxanthine and leads to AT-to-GC transitions after DNA replication (45). In *B. subtilis*, hypoxanthine has been studied both related to purine metabolism (38), and DNA damage and repair (46). To the best of our knowledge, this is the first time that hypoxanthine has been reported as a mediator of interactions in *B. subtilis* biofilms.

*L. fusiformis* is a free-living bacterium that can be isolated from soil and has been studied due to its production of interesting secondary metabolites and bioremediation potential (47–49). Here, we have identified a strain of *L. fusiformis* able to produce and excrete hypoxanthine in sufficient levels to induce the formation of wrinkles in biofilm colonies of *B. subtilis*. We found no evidence that this phenomenon is dependent on the signal transduction of a single Kin kinase (Fig. 5), and the expression levels of the matrix-component-related operons *epsA-O* and *tapA-sipW-tasA* remained equal when *B. subtilis* was exposed to the supernatant of *L. fusiformis* M5 (Fig. 6). Importantly, we cannot discard the possibility of changes in the expression of other genes. For example, changes in the expression of motility-related genes might be responsible for the apparent differences in colony expansion observed in our wrinkle formation assays. Another possibility is that an alternative exopolysaccharide biosynthetic pathway could be affected. Recently, *ydaJKLMN* was reported as a new operon important for the production of a so far unidentified exopolysaccharide in *B. subtilis* (50). However, overexpression of the *ydaK-N* operon under a xylose-inducible promoter could promote wrinkle formation in a Δ*epsH* mutant strain (50), while the presence of the *eps* operon is essential for the induction of wrinkle development in the presence of hypoxanthine. Therefore, it seems unlikely that hypoxanthine-induced wrinkle formation would proceed by directly inducing the production of an alternative exopolysaccharide. In contrast, we could detect the presence of dead cells with Sytox Green at the site of wrinkle formation (Fig. 8). Additionally, a knock-out mutant of the hypoxanthine/guanine permease PbuO lost the ability to form wrinkled colonies; specifically, a Δ*pbuG* mutant strain showed slightly reduced or similar wrinkle formation as *B. subtilis* 168, while a Δ*pbuO* strain completely lost the ability to form wrinkled colonies when exposed to hypoxanthine. Based on these results, we suggest that hypoxanthine induces the formation of wrinkles in colony biofilms of *B. subtilis* not by inducing the expression of biofilm-related genes, but rather by metabolic effects derived from the excess of available hypoxanthine. In this regard, we note that an excess of hypoxanthine can cause oxidative stress and cell death in eukaryotic cells by increasing the formation of reactive oxygen species when hypoxanthine is metabolized to urate (42, 43). A similar catabolic pathway could be followed by hypoxanthine in *B. subtilis,* which possesses multiple hypoxanthine/xanthine oxidases known as PucA, B, C, D and E (see SubtiWiki Pathways: http://www.subtiwiki.uni-goettingen.de/apps/pathway.php?pathway=41) (39). Additionally, it has been shown that localized cell death can lead to the formation of wrinkles in colonies of *B. subtilis* by providing an outlet for compressive mechanical forces that buckle the biofilm and promote the appearance of wrinkles (13). Thus, we hypothesize that the hypoxanthine provided by *L. fusiformis* induces oxidative stress and cell death in *B. subtilis*, which leads to the formation of wrinkles as a mechanical consequence. This is in accordance with the fact that the observed development of wrinkles only occurs in the interaction zone between colonies of these organisms (Fig. 1), or in the proximity of the cotton discs during our colony wrinkle formation assays (Figs. 5 and 9). This development of wrinkles does not happen in the rest of the *B. subtilis* colony, presumably due to a lower concentration of diffused hypoxanthine. Importantly, this induction of increased wrinkle formation in *B. subtilis* would therefore be a consequence of regular metabolic processes of *L. fusiformis*, rather than a canonical signaling mechanism intended to elicit a response in the receiver. Regarding hypoxanthine production by *L. fusiformis* M5, we have previously sequenced this strain, finding genes homologous to those known in other organisms to be responsible for hypoxanthine synthesis and export (51), although more research is needed to establish its production yield. However, we note that hypoxanthine can be produced by spontaneous deamination of adenine, such as that present in the surroundings of decaying cells (52). Unfortunately, our efforts to transform *L. fusiformis* M5 failed, preventing us to construct mutants with altered hypoxanthine production.

In recent years, biofilm research has grown from an incipient field to a major area of microbiological interest. Due to the high cell density of biofilms, social interactions are an inherent characteristic of these microbial populations, regardless of whether they are formed as single or multi-species communities (10, 28, 53, 54). The interactions between the organisms forming a biofilm are therefore an important aspect of this research field, since they shape the development of these communities, be it by intraspecies signaling, interspecies communications, or chemical cues derived from the metabolism of community members, such as the case presented here. Further study of the sociomicrobiology of biofilms will lead to an increased understanding of these communities as they form in nature, better enabling us to eliminate them when they are noxious to human activities, or to promote them when needed for biotechnological applications.

## MATERIAL AND METHODS

### Strains, media, and general culture conditions

All strains used in this study are listed in Table 1. When fresh cultures were needed, these strains were pre-grown overnight in Lysogeny broth medium (LB-Lennox, Carl Roth; 10 g L^−1^ tryptone, 5 g L^−1^ yeast extract, and 5 g L^−1^ NaCl) at 37°C and shaken at 225 r.p.m. LB medium was used for all *B. subtilis* and *Escherichia coli* transformations, and to screen soil samples. 2×SG medium (16 g L^−1^ nutrient broth (Difco), 2 g L^−1^ KCl, 0.5 g L^−1^ MgSO_4_·7H_2_O, 1 mM Ca(NO_3_)_2_, 0.1 mM MnCl_2_·4H_2_O, 1 µM FeSO_4_, and 0.1% glucose) (55) was used to grow cultures intended for supernatant production. This medium was also used for all strain interaction assays and wrinkle-induction assays. Tryptone Soya broth (CASO-Bouillon, AppliChem; 2.5 g L^−1^ glucose, 5 g L^−1^ NaCl, 2.5 g L^−1^ buffers (pH 7.3), 3 g L^−1^ soya peptone, and 17 g L^−1^ tryptone) was used for screening soil samples. GCHE medium (1% glucose, 0.2% glutamate, 100 mM potassium phosphate buffer (pH: 7), 3 mM trisodium citrate, 3 mM MgSO_4_, 22 mg L^−1^ ferric ammonium citrate, 50 mg L^−1^ L-tryptophan, and 0.1% casein hydrolysate) was used to induce natural competence in *B. subtilis* (56). Our developed Gallegos Rich medium was used to grow *Lactococcus lactis* MG1363, in order to purify pMH66: 21 g L^−1^ tryptone, 5 g L^−1^ yeast extract, 8.3 g L^−1^ NaCl, 3 g L^−1^ soya peptone, 2.6 g L^−1^ glucose, and 2.5 g L^−1^ MgSO_4_·7H_2_O. Overnight cultures of *L. lactis* were incubated at 30°C without shaking. Media were supplemented with Bacto agar 1.5% when solid plates were needed. Antibiotics were used at the following final concentrations: kanamycin, 10 µg mL^−1^; chloramphenicol, 5 µg mL^−1^; erythromycin-lincomycin, 0.5 µg mL^−1^ and 12.5 µg mL^−1^ respectively; ampicillin, 100 µg mL^−1^; spectinomycin, 100 µg mL^−1^; tetracycline, 10 µg mL^−1^. Specific growth conditions are described in the corresponding methods section. Importantly, all 2×SG plates used in this study were prepared with 25 mL of medium, and dried for a minimum of 20 minutes before use. Insufficient drying resulted in excessive colony expansion without development of architecturally complex colonies. To dry the plates, they were first allowed to solidify at room temperature for 1 hour, afterwards, they were kept completely open in a laminar flow sterile bench for the duration of the drying period. These drying conditions were followed for all assays that examined changes in colony architecture.

**Table 1.** Strains and plasmids used in this study.

| Strain | Characteristics | Reference |
| --- | --- | --- |
| *B. subtilis* |  |  |
| 168 | 168 1A700 *trpC*. Jena Stock | (33) |
| TB48 | 168 *trpC2 amyE::P<sub>hyperspank</sub>-mKATE (cat)* | (61) |
| JH12638 | JH642 *trpC2 phe-1 kinA::Tn917(cat)* | (62) |
| JH19980 | JH642 *trpC2 phe-1 kinB::tet* | (63) |
| RGP0203-4 | JH642 *kinC::Sp<sup>R</sup>* | (64) |
| DL153 | NCIB 3610 *kinD::tet* | (65) |
| BKE06370 | 168 *trpC2 pbuG::Erm<sup>R</sup>* | (66) |
| BKE13530 | 168 *trpC2 kinE::Erm<sup>R</sup>* | (66) |
| BKE29990 | 168 *trpC2 pbuO::Erm<sup>R</sup>* | (66) |
| NRS2243 | NCIB 3610 *sacA::P<sub>epsA</sub>-gfp (neo), hag::cat* | (67) |
| NRS2394 | NCIB 3610 *sacA::P<sub>tapA</sub>-gfp (neo)* | (67) |
| Δ*eps* | 168 *trpC2 epsA-O::tet* | (68) |
| Δ*tasA* | 168 *trpC2 tasA::Km<sup>R</sup>* | (68) |
| TB150 | 168 *trpC2 amyE::P<sub>hyperspank</sub>-mKATE (cat) epsA-O::tet* | This study,<br>TB48→Δ*eps* |
| TB171 | 168 *trpC2 amyE::P<sub>hyperspank</sub>-mKATE (cat) tasA::Km<sup>R</sup>* | This study,<br>TB48→Δ*tasA* |
| TB812 | 168 *trpC2 amyE::P<sub>hyperspank</sub>-mKATE (cat) pbuG::Erm<sup>R</sup>* | This study,<br>BKE06370→TB48 |
| TB813 | 168 *trpC2 amyE::P<sub>hyperspank</sub>-mKATE (cat) pbuO::Erm<sup>R</sup>* | This study,<br>BKE29990→TB48 |
| TB822 | 168 *trpC2 amyE::P<sub>hyperspank</sub>-mKATE (cat) Δ*pbuO** | This study |
| TB823 | 168 *trpC2 amyE::P<sub>hyperspank</sub>-mKATE (cat) Δ*pbuO, pbuG::Erm<sup>R</sup>** | This study,<br>BKE06370→TB822 |
| TB833 | 168 *trpC2 amyE::P<sub>hyperspank</sub>-mKATE (cat) kinA::Tn917(cat)* | This study,<br>JH12638→TB48 |
| TB834 | 168 *trpC2 amyE::P<sub>hyperspank</sub>-mKATE (cat) kinB::tet* | This study,<br>JH19980→TB48 |
| TB835 | 168 *trpC2 amyE::P<sub>hyperspank</sub>-mKATE (cat) kinC::Sp<sup>R</sup>* | This study,<br>RGP0203-4→TB48 |
| TB836 | 168 *trpC2 amyE::P<sub>hyperspank</sub>-mKATE (cat) kinD::tet* | This study,<br>DL153→TB48 |
| TB869 | 168 *trpC2 amyE::P<sub>hyperspank</sub>-mKATE (cat) sacA::P<sub>epsA</sub>-gfp (neo)* | This study,<br>NRS2243→TB48 |
| TB870 | 168 *trpC2 amyE::P<sub>hyperspank</sub>-mKATE (cat) sacA::P<sub>tapA</sub>-gfp (neo)* | This study,<br>NRS2394→TB48 |
| TB911 | 168 *trpC2 amyE::P<sub>hyperspank</sub>-mKATE (cat) kinE::Erm<sup>R</sup>* | This study,<br>BKE013530→TB48 |
| *E. coli* |  |  |
| MC1061 | Cloning host; K-12 F<sup>-</sup> $\lambda^-$ $\Delta(ara-leu)$7697 [araD139]B/r $\Delta(codB-lacI)$3 galK16 galE15 e14<sup>-</sup> mcrA0 relA1 rpsL150(Str<sup>R</sup>) spoT1 mcrB1 hsdR2(r<sup>-</sup> m<sup>+</sup>) | (69) |
| Soil Isolates |  |  |
| *Lysinibacillus* sp. M2c |  | This study |
| *Lysinibacillus fusiformis* M5 |  | This study |
| *Bacillus pumilus* P22a |  | This study |
| *Acinetobacter variabilis* T7a |  | This study |
| **Plasmid** | **Characteristics** | **Reference** |
| pMH66 | pNZ124-based Cre-encoding plasmid, Tet<sup>R</sup> Ts | (70) |

### Strain construction

All *B. subtilis* strains generated in this work were obtained via natural competence transformation (56) using genomic or plasmid DNA from donor strains as indicated in Table 1. Briefly, overnight cultures of the receiver strains were diluted to a 1:50 ratio with GCHE medium, these cultures were incubated at 37°C for 4 h with shaking at 225 r.p.m. After this incubation period, 5–10 µg of genomic or plasmid DNA were mixed with 500 µL of competent cells and further incubated for 2h before plating on LB plates added with selection antibiotics. Strain TB822 was obtained by using the Cre recombinase expressed from plasmid pMH66 to eliminate the Erm^R^ cassette of TB813, and subsequently curating pMH66 via thermal loss of the plasmid (57). Briefly, TB813 was transformed with 10 µg of pMH66, selecting transformants via incubation at 37°C on LB plates added with tetracycline. Candidates were then screened for their capacity to grow at 37°C on LB plates added with macrolide antibiotics (erythromycin-lincomycin), those that were not able to grow were further incubated on LB plates at 43°C for 18 h to induce the loss of pMH66. Candidates that were then unable to grow at 37°C on LB plates added with tetracycline were considered to have lost pMH66.

Successful construction of all used strains and plasmids was validated via PCR and restriction pattern analysis using standard molecular biology techniques, and by the lack of amylase activity on 1% starch LB plates (58) and emission of red fluorescence. All PCR primers used in this study are listed in Table 2. Primer pairs were used to amplify the indicated loci (see Table 2) in order to confirm the proper mutation of the corresponding gene. To confirm the correct construction of strains TB869 and TB870, primer oGFPrev2 was used in combination with oRGM38 (for *sacA*::P_*tapA*_-*gfp*) or oRGM40 (for *sacA*::P_*epsA*_-*gfp*).

**Table 2.**
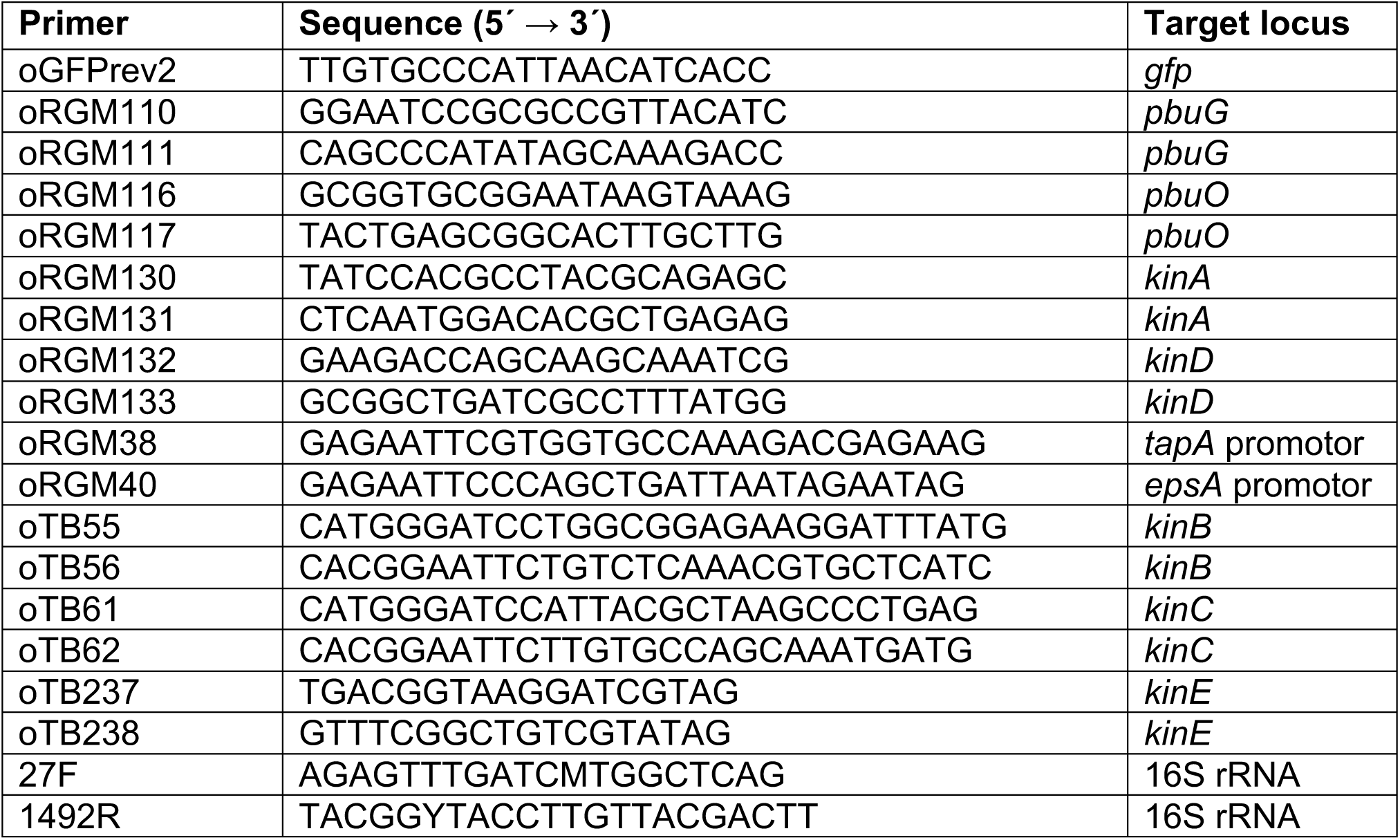
PCR primers used in this study.

### Isolation of bacteria from soil samples

Two independent Mexican soil samples were screened to isolate bacteria able to grow on LB or tryptone soya broth media. The first sample was collected from Tepoztlán, Morelos (18° 59′ 7″ N, 99° 5′ 59″ W), a humid and verdant region of central Mexico. The second sample was collected from Tehuacán, Puebla (18° 25′ 47.71″ N, 97° 27′ 58.1″ W), a semidesertic dry region in east central Mexico. Both samples were collected with a clean metal spatula 5 cm below surface level, and at 15 to 20 cm of the roots of local trees.

1 g of each soil sample was suspended in 9 mL of a sterile 0.85% NaCl solution, and 50 μL of Tween 80 were added. The resulting suspensions were vortexed for 5 minutes at maximum speed. The bigger soil particles were allowed to sediment by keeping the suspensions still for 10 minutes. The supernatants were then diluted with sterile 0.85% NaCl to 1:1000, 1:10 000 and 1:100 000 ratios. 50 and 100 μL of these dilutions were spread on LB and tryptone soya broth agar plates. These plates were incubated at 30°C for a maximum of 5 days. Bacterial colonies that grew during the incubation period were further isolated by cross-streaking them on LB or tryptone soya broth plates and incubating them at 30°C for 48 hours. Single isolated colonies obtained from this secondary cultivation were used to prepare liquid cultures on 3 mL of LB media. These cultures were incubated at 30°C with shaking for a maximum of 48 hours. Bacteria that grew efficiently during this incubation period were used to prepare glycerol stock solutions (20% v/v) and stored at −80°C for further use. In total, 242 soil strains were obtained and subsequently tested.

### Soil strains interaction screening

Overnight liquid cultures of *B. subtilis* strain TB48 and the obtained soil strains were adjusted to OD600 0.2 using LB medium. These diluted cultures were then mixed in 1:1, 10:1, and 1:10 ratios, and 2 µL of the mixed and pure cultures were inoculated on 2×SG plates. For neighbor colonies interaction assay, 2 µL of the pure cultures were inoculated at a distance of 5 mm from each other. Plates were incubated at 30°C for 72 h. The obtained colonies were used for microscopy analysis without further treatment.

### Identification of soil strains

Genomic DNA of selected soil strains was extracted with the GeneMATRIX Bacterial and Yeast Genomic DNA Purification Kit, according to the manufacturer’s instructions (EURx, Poland). This DNA was used to PCR amplify a fragment of the 16S rRNA gene using primers 27F and 1492R (59). Amplicons were purified with the High Pure PCR Product Purification Kit according to the manufacturer’s instructions (Roche Diagnostics, Switzerland) and sequenced with primers 27F and 1492R (GATC Biotech, Germany). Sequencing results were then compared with sequences in the National Center for Biotechnology Information (16S ribosomal RNA Bacteria and Archaea database) using the BLASTn algorithm (60). Soil strains’ identities were established using minimum query coverage of 98% and minimum identity values of 99%.

### Purification and treatment conditions of soil strain supernatant

To obtain cell-free supernatants of selected soil strains, 10 mL cultures on 2×SG medium of the corresponding strains were incubated at 30°C for 24 hours with shaking at 100 r.p.m. These cultures were then centrifuged at 7000 r.c.f. for 15 min. The supernatants were collected and filter-sterilized using a 0.22 µm pore-size filter (Carl Roth, Germany).

### Colony wrinkle formation assay

We developed the following assay to assess the effect that the supernatants of soil strains, and compound solutions, may have upon the architectural complexity of *B. subtilis* colony biofilms. Sterile 12 mm-diameter cotton discs were placed on 90 mm-diameter 2×SG agar plates, in such a way that there is a maximum amount of available space among the discs themselves and between the discs and the border of the plates. 3 cotton discs were used per plate. 50 µL of the tested supernatant or compound solution were deposited on the center of the cotton discs, and the plates were dried for 3 min by keeping them completely open in a laminar flow sterile bench. This drying period was done in addition to the regular 20 min drying previously described. 2 µL of an overnight culture of the tested strains were then inoculated at 5 mm from the edge of the cotton discs. The plates were incubated at 30°C for a total of 72 hours. Every 24 hours the cotton discs were reimpregnated with 25 µL of the corresponding supernatant or compound solution. After the incubation period, the plates were used for microscopy analysis without further treatment.

### Bioassay-guided fractionation

A 50 mL culture of M5 isolate was grown for 24 h under standard conditions. Bacterial cells were pelleted by centrifugation for 10 min at 6.000 r.c.f. and the supernatant was filtered through a 0.2 µm filter. 25 mL of the supernatant were freeze-dried and the remaining foam was dissolved in 1 mL water. The solution was applied to a Sephadex G20 column (3 cm × 40 cm) and eluted with water collecting 3 mL fractions (50 fractions). Fractions were further analyzed using the colony wrinkle formation assay. The fraction with the largest activity was analyzed using an LCMS (Shimadzu Deutschland, Germany) equipped with a Hypercarb column (100 × 3 mm, 3 µM, Thermo Fisher Scientific, flow rate = 0.6 mL min^−1^, method: 0-10 min: 100% (v/v) water). The main compound of this fraction was purified using a semi-preparative HPLC (Shimadzu Deutschland, Germany) equipped with Hypercarb column (100 × 10 mm, 5 µM, Thermo Fisher Scientific, flow rate = 5 mL min^−1^, method = 0-20 min: 100% (v/v) water). The pure compound was analyzed using ^1^H-NMR spectroscopy, LCMS, and HR-ESIMS. Obtained analytical data were in good agreement with a hypoxanthine standard from Sigma Aldrich.

### Comparison of hypoxanthine production

Supernatant of different isolates were heated to 80°C for 15 min and filtered through a 0.2 µm syringe filter. The samples were analyzed using an LCMS (Shimadzu Deutschland, Germany) equipped with a Hypercarb column (100 × 3 mm, 3 µM, Thermo Fisher Scientific, flow rate = 0.6 mL min^−1^, method: 0-10 min: 100% (v/v) water).

### Cell death assessment

To visualize cell death in colony biofilms we performed the colony wrinkle formation assay on plates supplemented with 0.25 µM Sytox Green nucleic acid stain (Thermo Fisher Scientific, U.S.A.). After 72 h of incubation, sectors of the colonies that grow around the cotton discs were manually sliced with a scalpel to produce thin cross-sections that include the supporting agar and a sliver of colony biofilm. The cross-sections were placed on a glass slide and used for microscopy without further treatment. All transmitted light and fluorescence images of colony biofilm cross-sections were obtained with an Axio Observer 780 Laser Scanning Confocal Microscope (Carl Zeiss, Germany) equipped with an EC Plan-Neofluar 10x/0.30 M27 objective, an argon laser for stimulation of fluorescence (excitation at 488 nm for green fluorescence and at 561 nm for red fluorescence, with emissions at 528/26 nm and 630/32 nm respectively), a halogen HAL-100 lamp for transmitted light microscopy and an AxioCam MRc color camera (Carl Zeiss).

### Stereomicroscopy and Image Analysis

All bright-field and fluorescence images of colonies were obtained with an Axio Zoom V16 stereomicroscope (Carl Zeiss, Germany) equipped with a Zeiss CL 9000 LED light source, a PlanApo Z 0.5× objective, HE 38 eGFP filter set (excitation at 470/40 nm and emission at 525/50 nm), HE 63 mRFP filter set (excitation at 572/25 nm and emission at 629/62 nm), and AxioCam MRm monochrome camera (Carl Zeiss, Germany).

Images were obtained with exposure times of 20 ms for bright-field, and 2500 ms for red fluorescence or 3000 ms for green fluorescence when needed. For clarity purposes, the images of colonies are presented here with adjusted contrast and the background has been removed, so that the colony structures can be easily appreciated. The modified pictures were not used for any fluorescence measurements.

To assess the expression levels of P_*epsA*_-*gfp*, P_*tapA*_-*gfp,* and P_*hyperspank*_-*mKATE* fluorescent reporter fusions in colonies we used ImageJ (National Institutes of Health, USA). Briefly, the average fluorescence emission intensities were measured in the green fluorescence channel for GFP and red fluorescence channel for mKATE by using a region of interest (ROI) that surrounds the cotton discs in the pictures as a partial ring, taking care to avoid the disc area itself. The ROI was drawn in such a way that it avoids the region of the colony that first makes contact with the cotton discs. This ROI had a width of 0.5 mm for measurements done at 40 and 50 hours, and a width of 1 mm for measurements done at 65 hours. The ROI was positioned in each colony image using the bright-field channel, and the average fluorescence intensity was then measured on the corresponding green and red fluorescence images.

To assess cell death, the average green fluorescence intensity of the cross-sections of colonies treated with Sytox Green was measured. All measurements were done with ImageJ. The colony area was selected on the transmitted light channel of cross-section images using the tracing tool (Legacy mode, tolerance 60). The average fluorescence intensity was then measured in the corresponding areas of the green and red fluorescence channels.

### Nucleotide sequence accession numbers

Sequences used in this study have been deposited in GenBank under accession numbers KY705015 (*Lysinibacillus* sp. M2c), KY698015 (*Bacillus pumilus* P22a), and KY703395 (*Acinetobacter variabilis* T7a). Further, the draft genome sequence of *L. fusiformis* M5 is available in GenBank under accession number MECQ00000000, and the strain was deposited in the Jena Microbial Resource Collection (ST-Number: ST036146, see http://www.leibniz-hki.de/en/jena-microbial-resource-collection.html).

## AUTHORS CONTRIBUTIONS

R.G.-M., Á.T.K. conceived the project; R.G.-M., S.K. performed the screen and the microbiology assays; R.G.-M. constructed bacterial strains; S.G., purified hypoxanthine; S.G., R.B., P.S. analyzed the analytical data; R.G.-M., P.S., Á.T.K., wrote and revised the manuscript.

## ACKNOWLEDGEMENTS

We would like to thank Heike Heineke (HKI Jena) for NMR measurements and Andrea Perner (HKI Jena) for HR-MS measurements.

Ákos T. Kovács was funded by a Marie Curie Career Integration Grant (PheHetBacBiofilm), Deutsche Forschungsgemeinschaft (DFG) (KO4741/2-1 and KO4741/3-1), and DFG Graduate School Jena School of Microbial Communication (JSMC) (Start-up Grant). Ramses Gallegos-Monterrosa was funded by Consejo Nacional de Ciencia y Tecnología, German Academic Exchange Service. Sebastian Götze, Robert Barnett, and Pierre Stallforth were funded by Leibniz Association (Junior Research Group of Pierre Stallforth). Robert Barnett was funded by DFG Graduate School Jena School of Microbial Communication (JSMC) (PhD fellowship to Robert Barnett).

